# SAMSA: A comprehensive metatranscriptome analysis pipeline

**DOI:** 10.1101/046201

**Authors:** Samuel T Westreich, Ian Korf, David A. Mills, Danielle G Lemay

**Author notes:** Email addresses: STW IK DAM DGL.

## Abstract

**Background:** Although metatranscriptomics—the study of diverse microbial population activity based on RNA-seq data—is rapidly growing in popularity, there are limited options for biologists to analyze this type of data. Current approaches for processing metatranscriptomes rely on restricted databases and a dedicated computing cluster, or metagenome-based approaches that have not been fully evaluated for processing metatranscriptomic datasets. We created a new bioinformatics pipeline, SAMSA, designed specifically for metatranscriptome dataset analysis, which runs either inhouse or in conjunction with Metagenome-RAST (MG-RAST) servers. Designed for use by researchers with relatively little bioinformatics experience, SAMSA offers a breakdown of metatranscriptome activity by organism or transcript function, and is fully open source. We next used this new tool to evaluate best practices for sequencing stool metatranscriptomes.

**Results:** Working with the MG-RAST annotation server, we constructed the Simple Annotation of Metatranscriptomes by Sequence Analysis (SAMSA) software package, a complete pipeline for the analysis of gut microbiome data. In creating this package, we determined optimal parameters in data collection and processing. SAMSA can summarize and evaluate raw annotation results, identifying abundant species and significant functional differences between metatranscriptomes.

Using pilot data and simulated subsets, we determined experimental requirements for fecal gut metatranscriptomes. Sequences need to be either long reads (longer than 100bp) or paired-end reads that can be joined. Each sample nees 40-50 million raw sequences which can be expected to yield the 5-10 million annotated reads necessary for accurate abundance measures. We also demonstrated that ribosomal RNA depletion does not equally deplete ribosomes from all species within a sample, and remaining rRNA sequences should be discarded. Using publicly available metatranscriptome data in which rRNA was not depleted, we were able to demonstrate that organism activity can be measured using mRNA counts. We were also able to detect significant differences between control and experimental groups in both organism activity and functional activity.

**Conclusions:** By making this new pipeline publicly available, we have created a powerful new tool for metatranscriptomics research, offering a new method for greater insight into the activity of diverse microbial communities. We further recommend that stool metatranscriptomes be ribodepleted and sequenced in a 100bp paired end format with a minimum of 40 million reads per sample.

## Background

Metatranscriptomics, the large-scale sequencing of mRNAs from complex microbial communities, allows for the observation of gene expression patterns [1–4]. Metatranscriptomics is a relatively new field, with the first mention around 2008[5], but it is growing quickly. Using high-throughput techniques developed in conjunction with “big data” computer-automated analysis approaches, metatranscriptomics offers a novel and complete method for looking at not just the organisms present, but also the activity occurring within a complex and diverse population at any chosen specific point in time[2, 3]. Metatranscriptomics is especially useful for analyzing complex populations in flux, such as the gut microbiome, which can be impacted and altered by a large number of transitory factors [6–8].

However, because of the complexity of metatranscriptomic data, extensive analysis is needed to convert raw data into simplified and easily understood results. The raw data comprise tens of millions of individual reads per sample[4]. Simplifying and condensing this very large data set requires multiple steps in a software pipeline and generally requires a dedicated bioinformatician to perform the analysis. Current pipelines or in-house methods often require significant computing power or use several different tools, many of which were not originally intended for metatranscriptome analysis [2, 3, 9]. For researchers who want to utilize metatranscriptomic analysis but may not have the necessary bioinformatics experience, there is a strong need for a complete pipeline, designed to analyze this data from beginning to end without requiring extensive technical expertise.

In addition, as metatranscriptomics is a new area of genomic exploration, there are few established guidelines or standard protocols. Standardized protocols exist for mRNA collection [5] and for stool-specific extraction [10, 11], but there is also a pressing need for an investigation of necessary sequencing parameters and depth, minimum read quality standards, and reliable reference databases against which metatranscriptome reads can be aligned. Any researcher planing to sequence metatranscriptomic data must know the target sequencing depth and minimum recommended read length, whether to filter out ribosomal sequences, and the relative confidence of their predicted results based on these parameters.

In this paper, to develop a standard metatranscriptomic analysis pipeline that can be applied to a wide range of microbiome samples, we extracted and sequenced RNA from human fecal material using experimentally validated protocols [10]. By testing various parameters and modifications, we determined the best practices at each step in obtaining and analyzing the metatranscriptomic data. In addition, we incorporated a publicly available metatranscriptomic data set examining the fecal gut microbiome in tyrosine kinase 2 knockout (Tyk2-/-) mice compared to wild-type controls over time during the outbreak of dextran sodium sulfate (DSS) induced colitis [12]. Examining this data, we provide new insights into differences in functional expression between the two different microbiomes over the course of the colitis development. By applying the approaches detailed here, researchers wishing to include metatranscriptomic analysis in their experiments can obtain accurate results without an investment in the development of bioinformatics tools.

## Results

### Creation of SAMSA, a new pipeline for metatranscriptome analysis

We created the SAMSA (Simple Analysis of Metatranscriptome Sequence Annotations) pipeline, designed to fully analyze and characterize activity within a metatranscriptome, illustrating relative level of activity split by both organism and functional category. There are four phases to the pipeline: the preprocessing phase trims and combines reads for input to the annotation phase, the annotation phase provides an annotation for each read, the aggregation phase aggregates organsim and function information across all reads, and the analysis phase provides visualizations and statistical analysis (sFigure 1).

**Figure 1.**
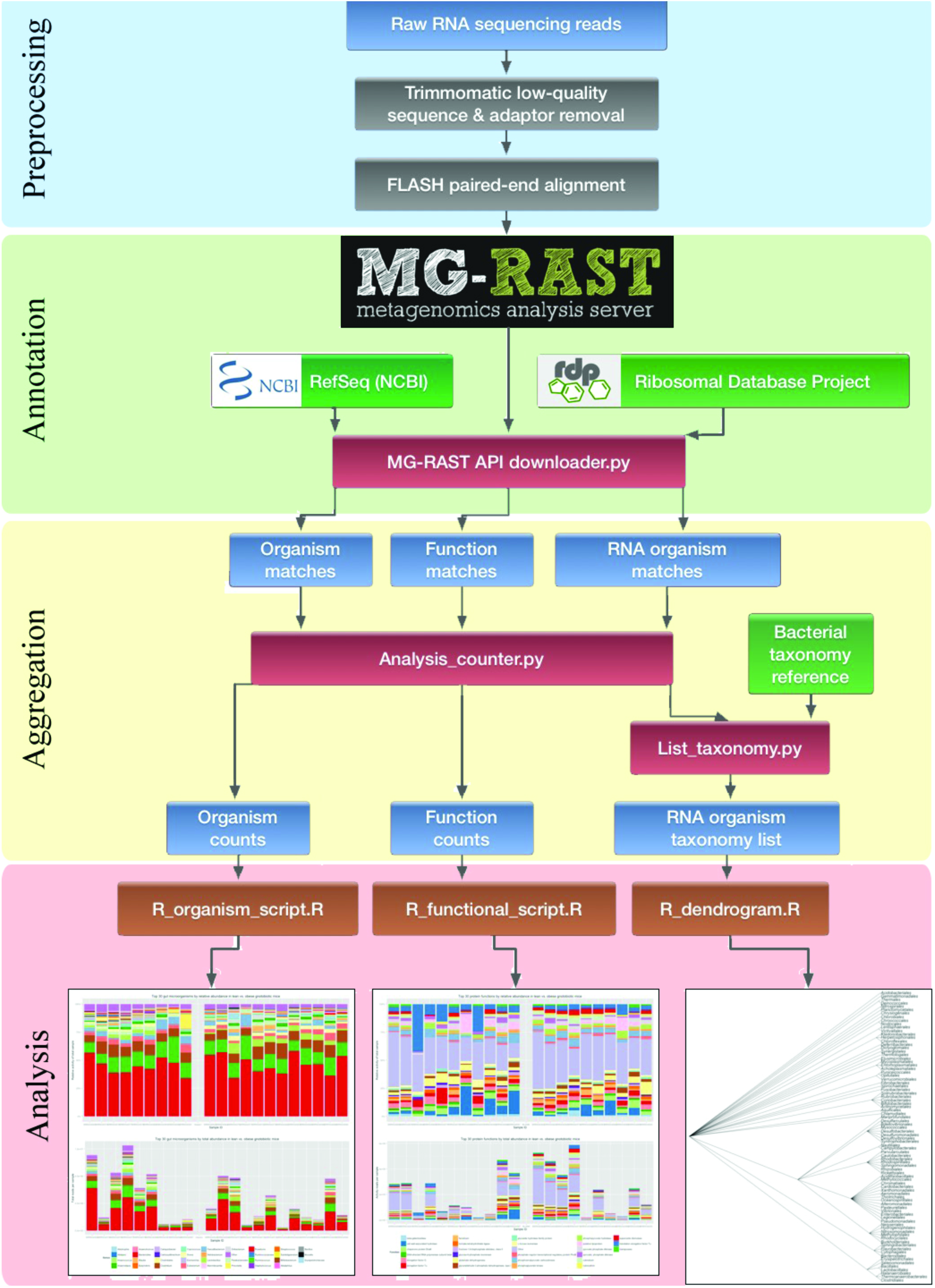
- the SAMSA pipeline. This organizational chart shows the flow of data through the pipeline, beginning with raw reads at the top of the chart and ending with the graphical output of the results at the bottom. Note that blue boxes denote intermediate generated output files, red boxes denote Python scripts, orange boxes denote R scripts, and green boxes denote external reference databases.

#### Preprocessing phase

During preprocessing, raw sequences are trimmed to remove reads containing low-quality bases and eliminate adaptor contamination using Trimmomatic, a flexible read trimming tool for Illumina NGS data[13]. Next, each pair of paired-end reads are aligned to each other using FLASh, a short read aligning program [14]. In our pilot samples, approximately 32-54% of the raw reads in each sample were successfully aligned, with an average aligned read length of 178 base pairs.

#### Annotation phase

Next, these sequences are submitted for annotation to Metagenomic Rapid Annotations using Subsystems Technology (MG-RAST) [15]. MG-RAST includes several steps, including an initial sequence quality control check through SolexaQA, gene calling through FragGeneScan, clustering of amino acid sequences at 90% identity through the uclust implementation of QIIME, and then using sBLAT on each protein sequence cluster to locate the best match reference. For each sequence cluster, MG-RAST selects the best match through the sBLAT similarity search. If multiple reference database matches tie for best matching score, they are both included in the final results. If the read does not achieve a match score above the minimum e-value cutoff, it is discarded. Each match is linked to MG-RAST’s internal identifier system and assigned an M5nr ID, correlating with linked matches in all subsystems databases. The annotated output can be provided on a per-match basis, using the M5nr ID to link each read to its best match from the subsystems database of choice. To create sorted abundance measures of the metatranscriptome using the SAMSA pipeline, all annotations with an acceptable best-match to the NCBI Reference Database (RefSeq) [16] are downloaded from MGRAST. Annotations are downloaded for the best match to both organism and individual transcript. Annotations are downloaded directly in tab-delimited form using MG-RAST’s RESTful API interface [17] and a custom Python program to assemble the API call command. In addition, the annotated output is also downloaded from the SEED Subsystems reference database [18] to provide ontology annotations.

#### Aggregation phase

A custom Python program parses each annotated output, storing each unique annotation match in a dictionary and maintaining counts of the number of occurrences of each unique annotation. After the annotation file is processed, unique annotations are sorted by abundance and exported as output.

#### Analysis phase

Annotation and abundance information from the Aggregation phase are inputs to custom R scripts, which generate barplots and dendograms (see Figure 1). To compare experimental versus control metatranscriptomes and determine significantly differentially expressed transcripts, R’s DESeq2 package is used to test the output files for differential expression [19]. DESeq2’s testing method adjusts for multiple hypothesis testing, performing pairwise comparisons in the scope of a larger overall data set. The pipeline’s R scripts export sorted lists of log2fold change in both organism and functional category activity, sorted by adjusted p-value. Results are also stored in the local R environment and can be used for graphical output.

This complete pipeline, coded in Python and R, is fully open source, is set up for streamlined use from the command line, does not require a local server for intense computation, and is freely available for download through GitHub.

### Measurement of host transcripts in fecal metatranscriptomes

Given that metatranscriptomes theoretically contain the transcriptional profile of all organisms present, we asked whether the transcriptomes of host cells could still be elucidated from fecal metatranscriptomes for which the primary goal was bacterial metatranscriptome analysis. We tried two methods of RNA extraction followed by either ribosomal depletion or poly(A) enrichment (see Methods). Neither method yielded many reads aligning to the human genome.We found 417 and 340 ribodepleted reads (0.0019% and 0.0085% of all ribodepleted reads) matching the human genome, and 2273 and 2100 poly(A) enriched reads (0.116% and 0.097% of all poly(A) enriched reads) matching the human genome. Thus, we conclude that host transcriptomes cannot be reliably extracted from stool metatranscriptomes that have been isolated and sequenced using standard methods.

### Use of ribosomal depletion methods in metatranscriptome preparation

Transcriptomic and metatranscriptomic extractions are often treated with ribodepletion methods before sequencing for the removal of ribosomal reads, increasing the relative mRNA yield [4, 20]. Comparing ribodepleted and non-depleted RNA, we found that the Ribo-Zero Gold ribodepletion kit from Illumina showed an overall 63-82% reduction in ribosomal reads present within our sample, but certain microbial ribosomes did not show any reduction, including members of the Actinobacteria, Cyanobacteria, and Spirochaetes phyla (Figure 2, Table 1). Ribodepletion appears effective at depleting rRNA reads within a sample, but also preferentially removes ribosomes from certain bacterial phyla over others, skewing ribosomal output within the metatranscriptome’s sequencing results.

**Figure 2.**
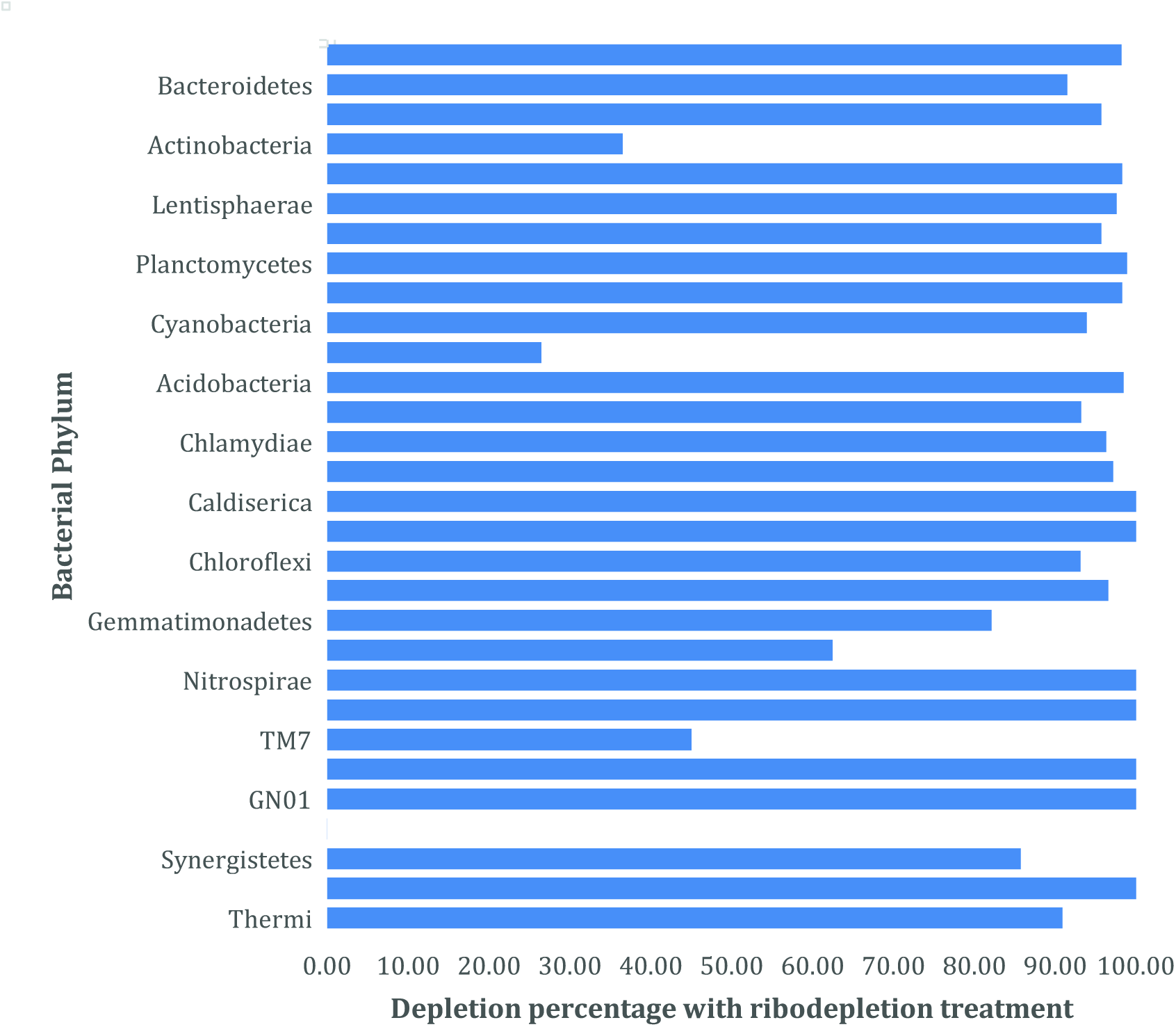
- Level of depletion in ribodepleted vs. control metatranscriptomes. Identical pilot metatranscriptome samples were sequenced; protocol 1 included a ribodepletion step, while protocol 4 did not include this step. As is demonstrated, not all species were equally depleted, skewing the perceived abundances of different organisms within the metatranscriptome. Normalized data shows the combined average depletion percentage for each phylum of bacteria present within the sample.

**Table 1.** - Comparison of rRNA abundance in control vs. ribodepleted samples. Comparisons of the organism annotations reveal that ribodepletion led to a decrease in 16S rRNA reads for most, but not all, bacterial phyla.

| **Phylum** | **4012, P4** | **4012, P1** | **4015, P4** | **4015, P1** | **% depleted** |
| --- | --- | --- | --- | --- | --- |
| Gemmatimonadetes | 3 | 1 | 28 | 5 | 74.40 |
| Spirochaetes | 3 | 4 | 14 | 55 | 163.10 |
| Nitrospirae | 4 | 1 | 22 | 0 | 87.50 |
| Fusobacteria | 5 | 5 | 24 | 9 | 31.25 |
| Fibrobacteres | 6 | 0 | 21 | 0 | 100.00 |
| Synergistetes | 6 | 5 | 14 | 2 | 51.19 |
| Thermi | 6 | 1 | 11 | 1 | 87.12 |
| Chlorobi | 6 | 1 | 34 | 0 | 91.67 |
| Chloroflexi | 7 | 6 | 29 | 2 | 53.69 |
| Caldiserica | 8 | 0 | 64 | 0 | 100.00 |
| Caldithrix | 11 | 0 | 70 | 2 | 98.57 |
| Tenericutes | 67 | 6 | 411 | 28 | 92.12 |
| Chlamydiae | 74 | 1 | 410 | 15 | 97.50 |
| Acidobacteria | 94 | 5 | 453 | 7 | 96.57 |
| Euryarchaeota | 175 | 15 | 942 | 693 | 58.93 |
| Parvarchaeota | 241 | 8 | 1298 | 22 | 97.49 |
| Cyanobacteria | 262 | 210 | 974 | 59 | 56.89 |
| Planctomycetes | 303 | 7 | 2019 | 23 | 98.28 |
| Crenarchaeota | 514 | 183 | 2474 | 106 | 80.06 |
| Lentisphaerae | 1045 | 19 | 6324 | 151 | 97.90 |
| Verrucomicrobia | 1107 | 29 | 6679 | 117 | 97.81 |
| Actinobacteria | 2739 | 5432 | 8394 | 5326 | 30.89 |
| Proteobacteria | 43585 | 3780 | 61436 | 2612 | 93.54 |
| Bacteroidetes | 486595 | 218584 | 1163164 | 98778 | 73.29 |
| Firmicutes | 939246 | 127130 | 2178859 | 38923 | 92.34 |
| **TOTALS** | **1476112** | **355433** |  |  | **68.57** |

### Optimal sequencing depth for gut microbiome metatranscriptomics

An important consideration when creating a metatranscriptome is sequencing depth: how many reads must be obtained in order to provide proper representation of all reads within a sample? As the proportionality of different reads within the sample is of crucial importance, sequencing depth must be large enough to ensure a balanced representation of each read’s relative abundance within the total sample. Too little depth can result in inaccurate abundance and activity measurements.

To evaluate necessary sequencing depth using a bioinformatics based approach, we generated 100 randomly selected subsets of a large and comparatively over-sequenced metatranscriptome of 21.6 million annotated reads (derived from 38M raw reads), creating ten smaller stand-alone simulated metatranscriptomes for each size point measured, moving in ten percent increments from 1 million up to 20 annotated million reads per subset. Within each simulated metatranscriptome, the relative percentage of various reads was measured, and these results were compared by simulated metatranscriptome size. This approach was performed looking at both high (in the 90^th^ percentile of all reads by sorted abundance), medium (in the 50^th^ percentile of all reads by sorted abundance), and low (in the 10^th^ percentile of all reads by sorted abundance) abundance reads within the parent metatranscriptome (Figure 3). By comparing relative abundance in each subset metatranscriptome to the final full data, we quantified the accuracy of the abundance measurements. We defined accuracy as percentage deviation from final, stable abundance counts. This data shows that, for the human gut metatranscriptomes examined, a minimum a minimum of five and ten million read annotations are needed to achieve above 90% accuracy in low abundance reads. This is equivalent to about 40-50 million raw sequence reads. Note that abundance estimates for medium and high abundance reads can be reasonably accurate with fewer annotations (Figure 3).

**Figure 3.**
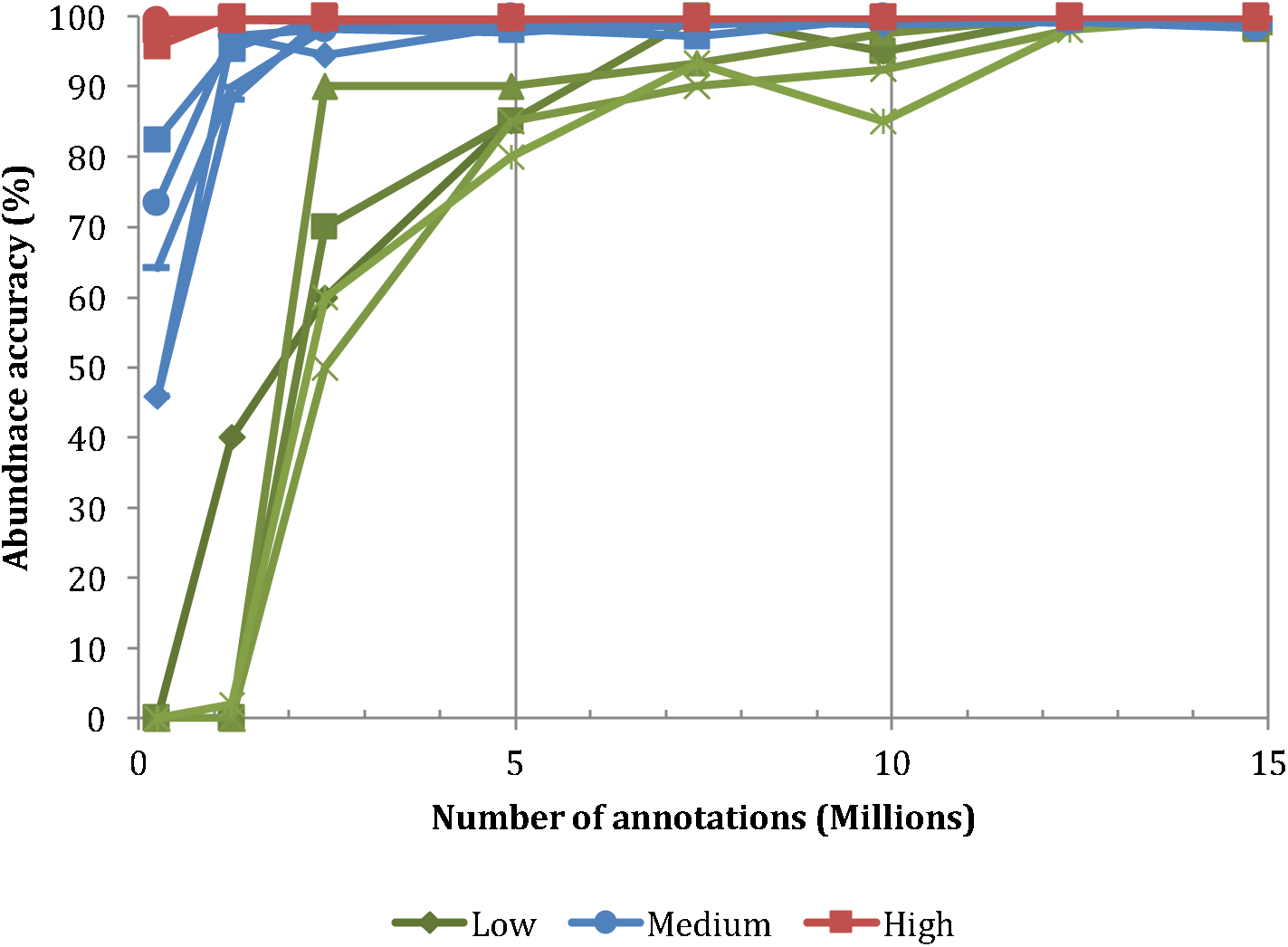
- Effects of metatranscriptome size on read abundance variation. As the number of annotations in a metatranscriptome increases along the X axis, accuracy of abundance measurements increases for all reads. Red denotes the top 5 most abundant transcripts within the sampled metatranscriptome (by counts), while blue denotes transcripts of medium abundance (top 50% by sorted counts) and green denotes low-abundance transcripts (bottom 10% by sorted counts). Approximately 10-15 million annotations are needed before abundance accuracy for all transcripts tops 90%. Abundance accuracy was measured out to 20 million annotations, but the accuracy was 100% in all categories beyond 15 million (data not shown).

### Paired or single end sequencing for metatranscriptome analysis

Another important consideration when sequencing a metatranscriptome is whether paired-end sequencing is necessary for proper read annotation. Paired-end sequencing is more expensive on a per-read basis but the average output read length is significantly increased, allowing for a more accurate best-match annotation in the pipeline.

Statistical comparisons of the relative organism activity results and the relative transcript abundance between single and paired end read versions of the same original metatranscriptome file show significant differences in both organism-of-origin and transcript identification by the annotation pipeline. In identical metatranscriptomes containing either paired-end or single end reads, the paired-end reads resulted in fewer reads removed due to quality controls (51,507 versus 123,518, or 0.3% vs. 0.6%), and a greater percentage of all reads matching at least one alignment in the reference database (77.2%, or 4,321,429 paired end reads versus 65.9%, or 3,194,911 single end reads) (Table 2).

**Table 2.** - Two pilot metatranscriptomes were sequenced twice; the P1 protocol involved a ribodepletion step, while the P4 protocol did not. Read count comparison between single and paired end files. Two pilot paired-end metatranscriptomes were analyzed both in paired configuration, and using only the forward reads (to simulate a single-end metatranscriptome from the same data). Despite higher numbers of total reads, the single-end data matched to fewer total and unique annotations in both cases.

|  | 4012, single read | 4012, paired end | 4015, single read | 4015, paired end |
| --- | --- | --- | --- | --- |
| Forward reads | 38,763,820 | - | 22,613,426 | - |
| Reverse reads | 38,763,820 | - | 22,613,426 | - |
| Paired assembled reads | - | 19,269,224 | - | 7,245,834 |
| Total annotations | 11,840,915 | 21,683,084 | 1,928,992 | 3,041,963 |
| Unique annotations | 8,658,502 | 13,310,506 | 1,280,940 | 1,875,267 |

The paired-end sequencing data matched to a higher number of unique transcripts than the single read data (270,384 unique transcripts versus 215,599 transcripts), suggesting that increased read length leads to greater specificity in transcript annotation. Correlations between paired-end and single-read processed identical metatranscriptomes averaged only 0.72 between both pilot samples. Given that these paired-end and single read metatranscriptomes were originally identical before the trimming of excess bases beyond 100 bases per read, these results indicate significant mislabeling or lost information in the single read approach when compared to the more accurate paired-end sample.

Taken with the organism matching results, these numbers suggest that the increased read length generated either through paired-end sequencing or through 150-bp single-end sequencing is necessary to ensure accurate metatranscriptome annotation.

### Determining organisms present within fecal metatranscriptomes

Using publicly available metatranscriptomes that were not depleted of ribosomal RNAs [12], we compared estimates of organism abundance using either mRNA or rRNA transcripts, using matches in NCBI’s RefSeq database for mRNA and matches in the SILVA small subunit (SSU) database for rRNA. We found a consistent correlation between abundance estimates from mRNA vs. rRNA (Figure 4) across all samples, suggesting that mRNA abundance estimates may be able to provide useful representative population data within a sample. With filtering to remove non-bacterial annotations, we observed an average Pearson correlation of 0.99 at the order level, 0.85 at the family level, and 0.81 at the genus level of identification across all 15 samples.

**Figure 4.**
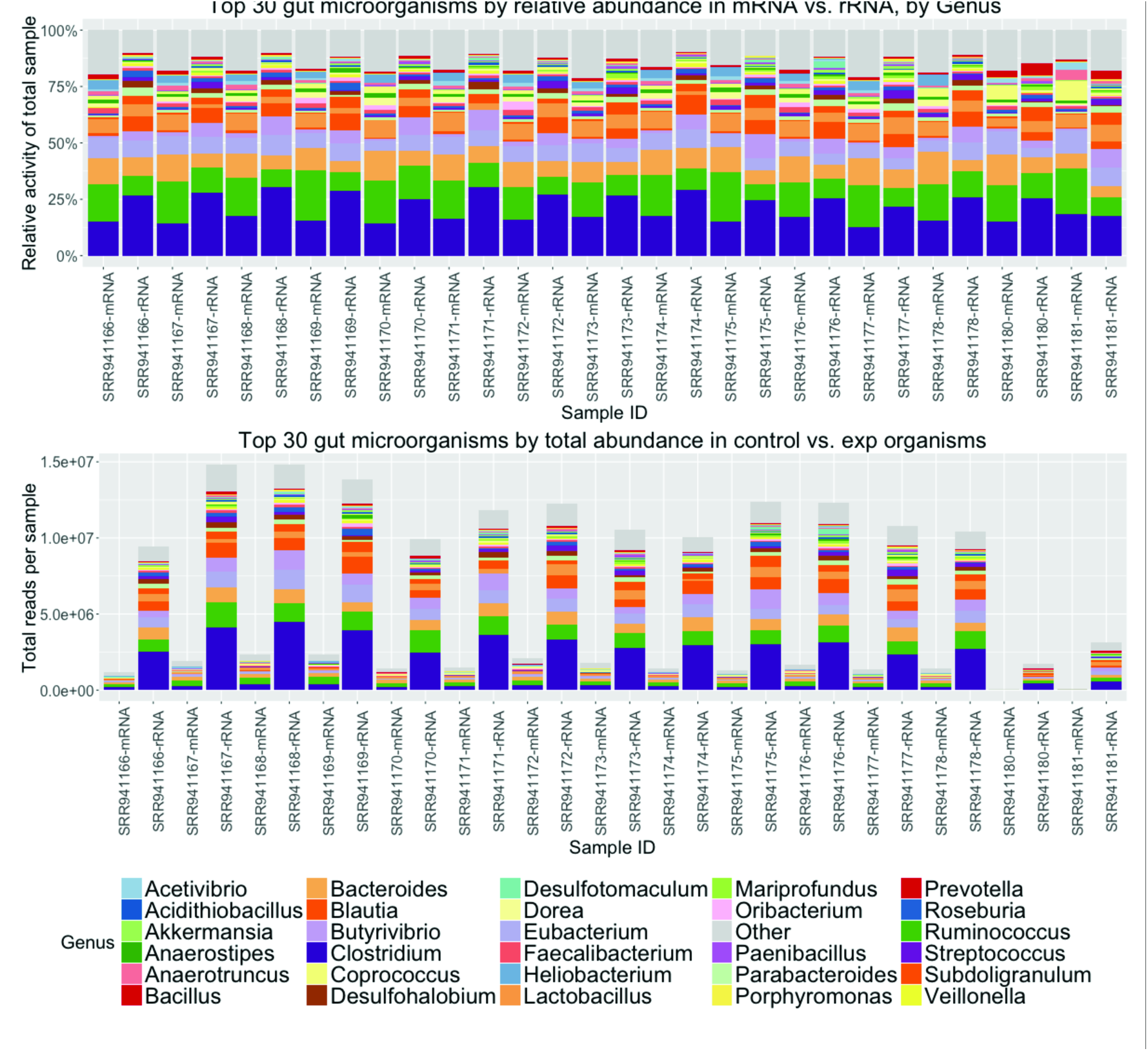
- Comparison of mRNA vs. rRNA based abundance estimates. A) Stacked bar graph measuring percentage distribution of total metatranscriptome activity by organism, with pairwise comparisons between mRNA and rRNA transcripts. To reduce potential mislabeling of organisms in the “long tail” of low-abundance organisms, only the top 30 most abundant organisms are displayed; other results are included in the purple “Other” catchall category. B) The same measurement, expressed in total number of annotations per metatranscriptome sample. Due to a lack of ribodepletion, rRNA transcripts dominate all samples.

### Applying this pipeline to a public metatranscriptomic data set: organism and functional analysis

Publicly available metatranscriptomic data [12] was analyzed using the SAMSA pipeline described above (Figure 1). Output tables generated by the pipeline are imported into R, where a stacked bar graph allows for comparing of relative activity levels both by organism (Figure 5) and by functional category (Figure 6). Due to the large number of both organism and functional categories, the R analysis scripts generate graphs which show only the top 30 most abundant organisms and/or functional categories, with remaining categories grouped under an “other” catch-all category (Figure 6A) or removed from display (Figure 6B).

**Figure 5.**
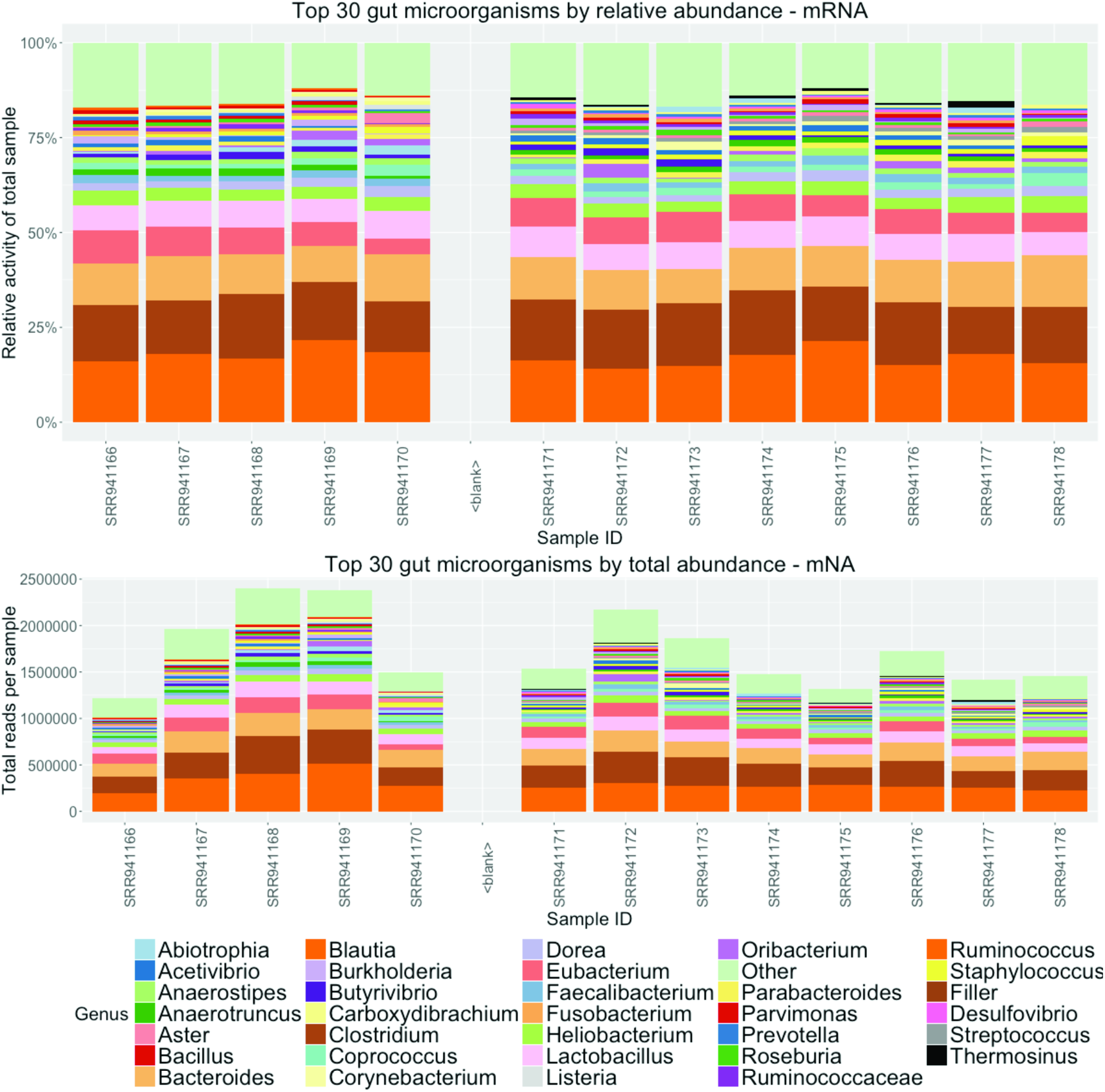
- Organism output of 15 metatranscriptomes. A) Stacked bar graph measuring percentage distribution of total metatranscriptome activity by organism, comparing 6 wildtype to 9 tyrosine kinase 2 knockout gut microbiomes. To reduce potential mislabeling of organisms in the “long tail” of low-abundance organisms, only the top 30 most abundant organisms are displayed; other results are included in the purple “Other” catchall category. B) The same measurement, expressed in total number of annotations per metatranscriptome sample.

**Figure 6.**
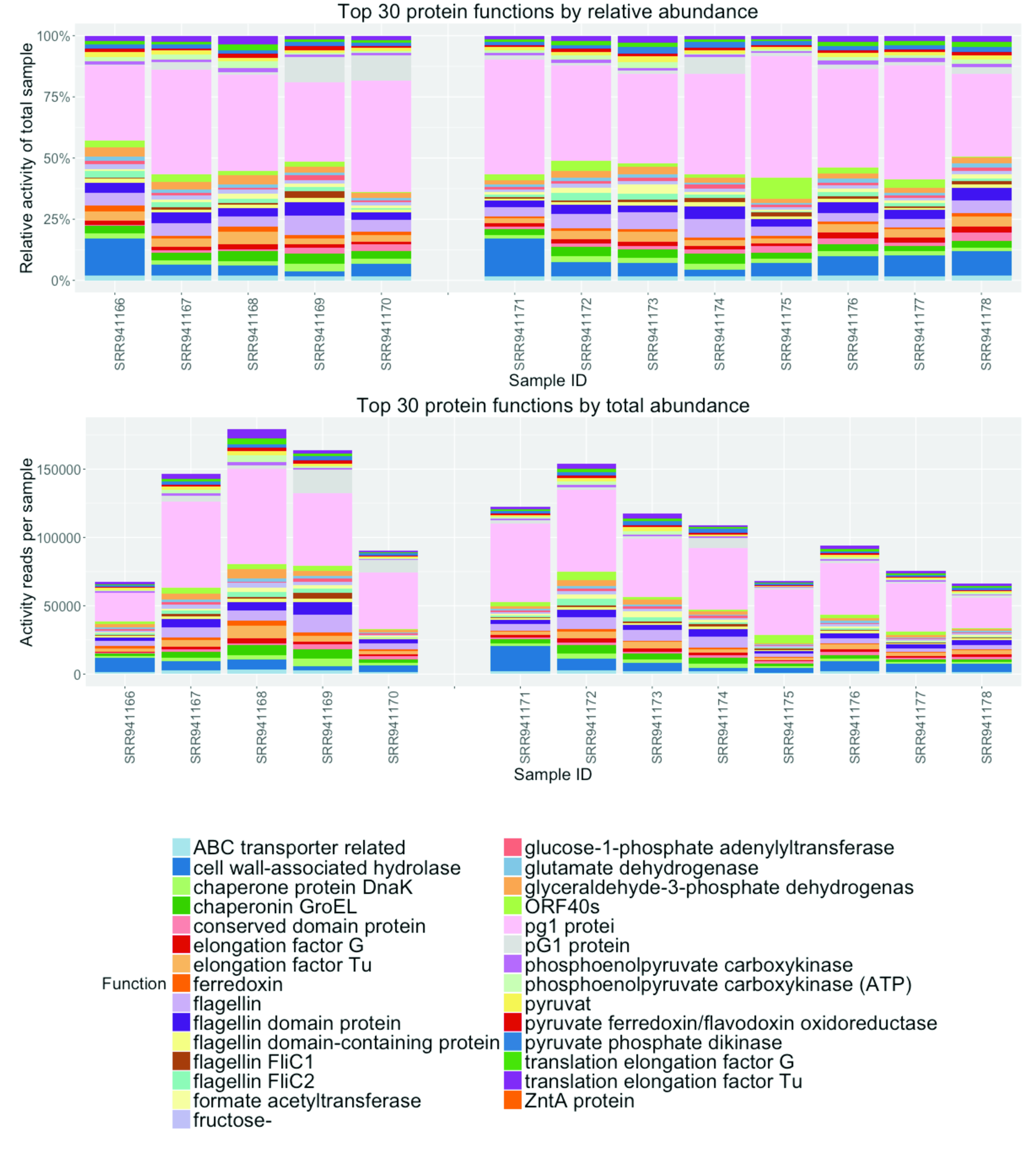
- Transcript functional category output of 13 metatranscriptomes. A) Stacked bar graph measuring percentage distribution of total metatranscriptome activity by functional category, comparing 5 wild-type to 8 tyrosine kinase 2 knockout mouse gut microbiomes. To reduce graph complexity, only the top 30 most abundant functions in each sample are displayed. Two samples were excluded due to very low total read counts. B) The same measurements, expressed in total number of annotations per metatranscriptome.

In the original paper [12], which focused primarily on the response of mouse gut epithelial cells to dextran sodium sulfate (DSS) induced colitis, the authors used primarily only the rRNA sequences from 16S amplicon sequencing of the samples to examine shifts in mouse gut microbiomes. They did report an increase in mRNA matching *Enterobacteriaceae* species in Tyrosine Kinase 2 (Tyk2) deficient mice, rising on day 3 of treatment, but provided no other mRNA analysis of these samples.

Examining the results of our pipeline, we observed a 1.558-fold increase in *Enterobacteriaceae* mRNA expression in day 3 samples (p=0.0055). Indeed, looking across all bacterial species with greater than 100 average expressed transcripts, we found significant increases in transcriptional activity in Escherichia and Providencia species, both of which are associated with ulcerative colitis [21]. In addition, we noted a significant decrease in Butyrivibrio species (p=0.03), when comparing between wild-type and Tyk2-deficient mice over the course of DSS-induced colitis. Butyrivibrio is a probiotic-associated strain that may normally offer protection against colitis [22]. We also observed several significant changes in functional activity between the two groups, identifying more than 300 protein-coding transcripts differentially expressed between wild-type and Tyk2-deficient mice (adjusted p<0.05, Additional File 1).

## Discussion

Although still an emerging field, metatranscriptomics offers a powerful approach for analyzing complex activity within a heterogeneous microbial population, such as that found within the gut microbiome. Previous approaches for the study of the gut microbiome provide an incomplete picture, limited to only identifying variations in organism population levels and not including data on the activity of different species within the environment.

Our pipeline fulfills the need for four main steps in metatranscriptomic data analysis: preprocessing, annotation, aggregation, and analysis. By leveraging MG-RAST’s annotation server, we can provide comprehensive annotation and analysis of a metatranscriptome without requiring a dedicated private server. SAMSA generates outputs at each step, creating a streamlined pipeline where each stage in data analysis can be independently examined.

Using the SAMSA pipeline, we have established a set of “best practices” for metatranscriptome sequencing. Our work demonstrates that the increased specificity provided by a paired-end sequencing approach significantly increases specificity of read annotations. In addition, to provide a complete and accurate measure of read abundance within a metatranscriptome, approximately forty million raw reads must be sequenced, providing an estimated ten million mRNA annotations.

Due to the vast majority of extracted total RNA originating from ribosomes, ribodepletion is strongly recommended for all metatranscriptome sample processing. We demonstrate that although ribodepletion is not successful at removing all rRNA, and skews the proportions of remaining rRNA within a sample, it greatly increases the number of mRNA annotations obtained per metatranscriptome. Although we only tested one commercially available ribodepletion kit, we believe that all ribodepletion methods based on hybridization with complementary rRNA oligonucleotide probes will carry some intrinsic bias, skewing the proportions of remaining rRNA within the sample. Therefore, we recommend discarding all remaining ribosomal reads before performing further analysis on these samples.

Although ribodepletion results in a loss of information regarding organism abundance within a metatranscriptomic sample, we show that total mRNA can be used as an alternate method of evaluating overall organism activity within a sample. Compared to mRNA results, we found a higher level of mismatches and false annotations among the rRNA data, particularly for eukaryotic organisms, including plant species such as Sisymbrium (cabbage), Arabidopsis, and Populus (cottonwood), Bos taurus, human, and Coptotermes (termite). This increased mismatch rate may be due to the fact that MG-RAST is not tailored to work with eukaryotic organisms.

Another reason that mRNA and rRNA results were not perfectly correlated is that there were insufficient mRNA reads in the data set used for direct comparison. Our data (Figure 3) suggested that a minimum of 10 million annotated reads are required for accurate annotations, while the non-ribosomal depleted data set averaged only 1-2 million annotated mRNA reads per sample. While total mRNA may not be a perfect proxy of organism activity, the extent of mismatches in the rRNA data suggest that mRNA is at least better than total rRNA for judging activity.

Analyzing a public data set, we confirmed the rRNA patterns previously described by the producers of that data [12], and also identified significant changes in other, more specific genera of microbial species, including increases in Escherichia and Providencia and decreases in Butyrivibrio. Both the increase in Escherichia and Providencia and the decrease in Butyrivibrio species were only notable at the genus level and would not be identifiable in 16S data, demonstrating the value of mRNA-based metatranscriptomics analysis. In addition, we successfully identified a variety of functional activities of gut microbes that significantly differed between wild-type and Tyk2-/- strains of mice strains.

As the field of metatranscriptomics continues to grow and expand, we expect that metatranscriptome analysis will become increasingly important to understand the functional responses of gut microbiome communities. By using mRNA transcripts to identify both the activity levels of organisms within a sample and changes in specific gene or functional expression, we can gain a better understanding of the capabilities and actions within an active gut microbiome at any chosen point in time. Because all parts of SAMSA are open source and publicly accessible, this tool can be used even by researchers with little previous experience in working with metatranscriptomes. We hope to encourage the more widespread use of metatranscriptomics as the next “big data” tool for determining activity within complex microbiome populations.

Although the SAMSA pipeline successfully annotated and summarized gut microbiome metatranscriptome data without the need for large server resources, several limitations still exist. MG-RAST’s annotation servers require waiting in a queue, and processing may be slowed by days to weeks. Additionally, although SAMSA can reveal shifts in activity patterns, shifts in microbial population sizes cannot be measured as accurately as using shotgun metagenomics, and future approaches may focus on incorporating both techniques to ensure completeness.

Future versions of this pipeline will focus on a more rapid in-house annotation step, using custom built reference databases and allowing for greater speed and additional analysis options, drawing from multiple reference databases as data progresses through the pipeline.

## Conclusions

We have created a new pipeline for metatranscriptome analysis, functioning in conjunction with the MG-RAST annotation pipeline. This pipeline is capable of determining expression activity within a sample at the transcript level, and provides measures of total activity, differentiated either by organism or by functional category of transcript. This pipeline will enable more rapid adoption of metatranscriptomics methods. Finally, we recommend that stool metatranscriptomes be ribodepleted and sequenced in a 100bp paired end format with a minimum of 40 million reads per sample.

## Methods

### Sample Collection

Stool samples were collected from two normal healthy adults. Stool was added to 50mL tubes containing 25mL RNAlater until the total volume reached 30mL. Samples were then sealed and shaked vigourously until the contents appeared to be uniform in consistency. The samples were then temporarily stored at −20 C before being moved to −80 C. These samples were derived from a study that is registered on clinicaltrials.gov (NCT01814540). The UC Davis Institutional Review Board approved all aspects of the study and written informed consent was obtained from all participants prior to study procedures.

### RNA extraction

Two different RNA extraction protocols were applied to the two human subject stool samples, adapted from Giannoukos et al.[10]. Samples were initially frozen at −80 degrees C in RNAlater. Samples were partially thawed and mixed with bacterial lysis buffer, incubated, and then homogenized through both bead-beating and QIAshredder treatments. Extraction was performed using the Qiagen RNeasy isolation kit, with additional rigorous Turbo DNAse treatment to remove DNA contamination. Ribodepletion was performed on two of the samples (designated P1) using the RiboZero Magnetic GOLD kit. Poly(A) selection was performed on two samples (designated P4) using the Illumina TruSeq kit protocol.

### RNA sequencing

For each sample, RNA-Seq libraries were prepared from 20uL of >2000ng/uL at the DNA Sequencing Core of the UC Davis Genome Center. RNA extracted using the bacterial metatranscriptome protocol (P1) was first ribodepleted using the RiboZero Magnetic Gold Kit (Epidemiology), catalog number MRZE706. The Illumina TruSeq protocol, without poly(A) selection, was then used to prepare RNA-Seq libraries. For the TRIzol-extract RNA (P4), the Illumina TruSeq protocol with poly(A) selection was used to prepare RNA-Seq libaries. All four samples were run on a single lane of Illumina HiSeq 2000 with indexing to allocate ~40% of the lane to each bacterial metatranscriptomes and ~10% of the lane to each poly(A)-selected metatranscriptome. The four metatranscriptomes have been deposited in the NCBI SRA repository, in BioProject PRJNA313102, SRA study SRP071017.

### Preprocessing and annotation of metatranscriptome reads

Raw sequences were obtained for two pilot samples, labeled as 4012 and 4015. Cleaning of the raw sequences to remove reads containing low-quality bases and eliminate adaptor contamination was performed using Trimmomatic, a flexible read trimming tool for Illumina NGS data. At default parameters, Trimmomatic removed low-quality reads to meet the minimum threshold of acceptability for MGRAST submission.

Paired end raw sequence files were aligned using FLASh, a short read aligning program. Approximately 32-54% of the raw reads in each sample were successfully aligned, with an average aligned read length of 178 base pairs.

The trimmed and aligned sequences were submitted for annotation to Metagenomic Rapid Annotations using Subsystems Technology (MG-RAST) [15]. MG-RAST includes several steps, including an initial sequence quality control check through SolexaQA, gene calling through FragGeneScan, clustering of amino acid sequences at 90% identity through the uclust implementation of QIIME, and then using sBLAT on each protein sequence cluster to locate the best match reference.

For each sequence cluster, MG-RAST selects the best match through the sBLAT similarity search. If multiple reference database matches tie for best matching score, they are both included in the final results. If the read does not achieve a match score above the minimum e-value cutoff, it is discarded. Each match is linked to MG-RAST’s internal identifier system and assigned an M5nr ID, correlating with linked matches in all subsystems databases. The annotated output can be provided on a per-match basis, using the M5nr ID to link each read to its best match from the subsystems database of choice.

### Post-annotation processing and analysis

To create sorted abundance measures of the metatranscriptome, all annotations with an acceptable best-match to the NCBI Reference Database (RefSeq) were downloaded from MG-RAST. Annotations were downloaded for the best match to both organism and individual transcript. Annotations were downloaded directly in tab-delimited form using MG-RAST’s RESTful API interface and a custom Python program to assemble the API call command. In addition, the annotated output was also downloaded from the SEED Subsystems reference database to provide ontology annotations.

A custom Python program parsed through each annotated output, storing each unique annotation match in a dictionary and maintaining counts of the number of occurrences of each unique annotation. After the annotation file was processed, the unique annotations were sorted by abundance and exported as output.

### Evaluating minimum viable metatranscriptome read counts

To determine minimum viable metatranscriptome size, the original samples, which were very deeply sequenced, were digitally broken down and reshuffled to create smaller, randomly generated subset metatranscriptomes. 100 smaller subset metatranscriptomes were generated for each subset size, with the subset sizes consisting of 1, 5, 10, 20, 30, 40, 50, 60, 70, 80, and 90 percent of the final metatranscriptome size.

Each of these smaller subsets was subjected to identical analysis using the same programs and pipeline to determine variation in annotation abundance. By comparing relative variation in the abundance of different transcripts, the minimum necessary metatranscriptome size needed for stable abundance percentage estimates could be computed.

To determine whether paired-end sequencing was necessary for ensuring accuracy of the annotation process, the paired-end reads were digitally trimmed to 100 base pairs, creating a simulated single-read metatranscriptome containing the same number of identical sequences to the paired-end file. This digitally created single-read metatranscriptome was analyzed using the same pipeline as the paired-end original file, and the results were compared for to determine level of variation using statistical testing in R via the DESeq2 package.

### Comparison of tyrosine kinase 2 knockout versus wild-type mice in colitis

The constructed pipeline was tested using publicly available RNA-seq data from Hainzl et al. [12]. The original authors obtained Illumina-sequenced metatranscriptomes from 15 gnotobiotic mice, 9 with a tyrosine kinase 2 knockout genotype (Tyk2-/-) and 6 wild-type controls, at different stages of dextran sodium sulfate (DSS) induced colitis. The original authors used only the rRNA gene copies from the metatranscriptomes, demonstrating through PCA analysis that both the wildtype and Tyk2-/- mice showed similar shifts in organism population based on stage of DSS-induced inflammation. Their raw data was made publicly available through NCBI’s Short Read Archive (SRA), accession numbers SRP026343 and SRP026649.

### Access to the SAMSA pipeline

All components and tools used in the SAMSA pipeline, as well as documentation files, are freely available from GitHub at http://github.com/transcript/SAMSA.

## Authors' contributions

DGL conceived of the study. DGL and STW designed experiments. STW developed the pipeline. STW and DGL conducted experiments. IK reviewed code. STW and DGL drafted the manuscript. STW, DGL, DAM, and IK interpreted data and contributed to the manuscript.

## Acknowledgements

The authors would like to thank Joyce Lee and Justin Fontaine for their technical help with RNA extraction from stool samples. We also thank Dr. Jennifer Smilowitz and Prof. Daniela Barile of University of California, Davis for their donation of the human stool samples. We also thank Folker Meyer and the team at MG-RAST for providing assistance with correcting API issues and inconsistencies. We thank Hainzl 2015 et al from the Institute of Animal Breeding & Genetics for making their data publically available through NCBI. This research was partially supported by an industry/campus supported fellowship of STW under the Training Program in Biomolecular Technology (T32-GM008799) at the University of California, Davis. The sequencing was carried out at the DNA Technologies and Expression Analysis Cores at the UC Davis Genome Center, supported by the NIH Shared Instrumentation Grant 1S10OD010786-01.

